# Molecular Mechanisms of Non-ionotropic NMDA Receptor Signaling

**DOI:** 10.1101/2020.01.08.898080

**Authors:** Ivar S. Stein, Deborah K. Park, Jennifer N. Jahncke, Juan C. Flores, Karen Zito

**Affiliations:** Center for Neuroscience, University of California, Davis, CA 95618

## Abstract

Structural plasticity of dendritic spines is a key component of the refinement of synaptic connections during learning. Recent studies highlight a novel role for the NMDA receptor (NMDAR), independent of ion flow, in driving spine shrinkage and LTD. Yet little is known about the molecular mechanisms that link conformational changes in the NMDAR to changes in spine size and synaptic strength. Here, using two-photon glutamate uncaging to induce plasticity in hippocampal CA1 neurons from mice and rats, we demonstrate that p38 MAPK is required downstream of conformational NMDAR signaling to drive both spine shrinkage and LTD at individual dendritic spines. In a series of pharmacological and molecular genetic experiments, we identify key components of the non-ionotropic NMDAR signaling pathway driving dendritic spine shrinkage, including the interaction between NOS1AP and nNOS, nNOS enzymatic activity, activation of MK2 and cofilin, and signaling through CaMKII. Our results represent a large step forward in delineating the molecular mechanisms of non-ionotropic NMDAR signaling that drive the shrinkage and elimination of dendritic spines during synaptic plasticity.

## Introduction

Activity-driven changes in neuronal connectivity are important for the experience-dependent remodeling of brain circuitry. In particular, the elimination of spine synapses is vital for the refinement of synaptic circuits throughout development and during learning. Indeed, an initial phase of spine formation and synaptogenesis during development is followed by a pruning phase leading to the removal of incorrect and redundant spine synapses (Wise et al., 1979; Holtmaat et al., 2005; Zuo et al., 2005). Furthermore, in vivo studies have shown that learning is associated with spine shrinkage and elimination, and that the level of spine loss is directly correlated with improved behavioral performance (Yang et al., 2009; Lai et al., 2012). Shrinkage and loss of dendritic spines is driven by glutamatergic signaling mechanisms leading to synaptic weakening through induction of long-term depression (LTD), and requires activation of the NMDA-type glutamate receptor (NMDAR) (Okamoto et al., 2004; Zhou et al., 2004; Hayama et al., 2013; Oh et al., 2013; Wiegert and Oertner, 2013).

Recent studies have demonstrated that NMDAR-dependent LTD and spine shrinkage can occur independent of ion flux through the NMDAR. Indeed, LTD and spine shrinkage induced by low frequency glutamatergic stimulation are blocked by competitive glutamate binding site NMDAR antagonists, but persist in the presence of the NMDAR glycine/D-serine binding site antagonist 7-CK or the pore blocker MK-801 ((Nabavi et al., 2013; Stein et al., 2015; Carter and Jahr, 2016; Wong and Gray, 2018), but see (Babiec et al., 2014)). Furthermore, high frequency glutamatergic stimulation that normally leads to LTP and spine growth instead drives LTD and spine shrinkage when ion flow through the NMDAR is blocked with 7-CK or MK-801 (Nabavi et al., 2013; Stein et al., 2015). Altogether, these findings support a model where glutamate binding triggers conformational changes in the NMDAR signaling complex, which, in the absence of Ca^2+^-influx, are sufficient to drive LTD and dendritic spine shrinkage.

Little is known about the molecular signaling mechanisms that link glutamate-induced conformational changes of the NMDAR to the induction of LTD and spine shrinkage. Non-ionotropic NMDAR signaling in LTD requires basal levels of intracellular Ca^2+^ and causes the activation of p38 MAPK (Nabavi et al., 2013). p38 MAPK is required for dendritic spine shrinkage induced by conformational signaling through the NMDAR (Stein et al., 2015). Furthermore, glutamate or NMDA binding causes conformational changes in the NMDAR intracellular domains that lead to changes in its interaction with the downstream signaling molecules PP1 and CaMKII, suggesting that these molecules could play an important role in NMDAR conformational signaling (Aow et al., 2015; Dore et al., 2015). Thus, while imaging experiments offer invaluable insights into the nature of the conformational and protein interaction changes, the only molecules directly implicated in non-ionotropic NMDAR signaling during synaptic plasticity to date are p38 MAPK and basal intracellular Ca^2+^.

Here we used two-photon glutamate uncaging, time-lapse imaging, and whole-cell recordings to define the molecular mechanisms that link non-ionotropic NMDAR signaling to the shrinkage and elimination of dendritic spines. We show that p38 MAPK is required for spine shrinkage driven by NMDAR conformational signaling not just in response to low frequency glutamatergic stimuli that induce LTD, but also in response to high frequency glutamatergic stimulation in the absence of ion flow through the NMDAR. We further found that this metabotropic NMDAR signaling drives synaptic weakening at individual dendritic spines, which also relies upon p38 MAPK, and that spine shrinkage does not require activation of AMPARs. Furthermore, we show that conformational NMDAR signaling in spine shrinkage relies on nNOS activation and on the interaction between nNOS and NOS1AP, linking p38 MAPK activation to the NMDAR signaling complex. Downstream of p38 MAPK, MAPK-activated protein kinase 2 (MK2) and cofilin are required to drive spine shrinkage. Finally, we show that spine shrinkage driven by conformational NMDAR signaling requires the activation of CaMKII. Our results delineate key components of the signaling pathway linking non-ionotropic NMDAR signaling to dendritic spine shrinkage.

## Material and Methods

### Preparation and transfection of organotypic slice cultures

Organotypic hippocampal slices were prepared from P6-P8 Sprague-Dawley rats or C57BL/6 mice of both sexes, as described (Stoppini et al., 1991). The cultures were transfected 1-2 d (EGFP alone) or 3-4 d (cofilin KD and rescue experiments) before imaging via biolistic gene transfer (180 psi), as previously described (Woods and Zito, 2008). We coated 6-8 mg of 1.6 µm gold beads with 10-15 µg of EGFP-N1 (Clontech) or 20 μg pSuper-cofilin1-shRNA + 20 μg pSuper-ADF-shRNA (Bosch et al., 2014) + 8 μg pCAG-CyRFP1 (Addgene; (Laviv et al., 2016)) + 4 μg EGFP-N1 or 20 μg pSuper-cofilin1-shRNA + 20 μg pSuper-ADF-shRNA + 8 μg pCAG-CyRFP1 + 4 μg shRNA insensitive cofilin1-EGFP (Bosch et al., 2014).

### Preparation of acute slices

Acute hippocampal slices were prepared from postnatal day 16-20 (P16-P20) GFP-M mice (Feng et al., 2000) of both sexes. Coronal 400 µm slices were cut (Leica VT100S vibratome) in cold choline chloride dissection solution containing (in mM): 110 choline chloride, 2.5 KCl, 25 NaHCO_3_, 0.5 CaCl_2_, 7 MgCl_2_, 1.3 NaH_2_PO_4_, 11.6 sodium ascorbate, 3.1 sodium pyruvate, and 25 glucose, saturated with 95% O_2_/5% CO_2_. Slices were recovered for 45 min in 30°C oxygenated artificial cerebrospinal fluid (ACSF) containing (in mM): 127 NaCl, 25 NaHCO_3_, 1.25 NaH_2_PO_4_, 2.5 KCl, 25 glucose, 2 CaCl_2_, and 1 MgCl_2_, and then incubated at room temperature for an additional 45 min before imaging.

### Time-lapse two-photon imaging

EGFP-transfected CA1 pyramidal neurons from acute (P16-P20) or cultured [14-18 days in vitro (DIV)] slices at depths of 10-50 µm were imaged using a custom two-photon microscope (Woods et al., 2011) controlled with ScanImage (Pologruto et al., 2003). Image stacks (512 × 512 pixels; 0.02 µm per pixel) with 1-μm z-steps were collected. For each neuron, one segment of secondary or tertiary basal dendrite was imaged at 5 min intervals at 29 °C in recirculating artificial cerebral spinal fluid (ACSF; in mM: 127 NaCl, 25 NaHCO_3_, 1.2 NaH_2_PO_4_, 2.5 KCl, 25 D-glucose, aerated with 95%O_2_/5%CO_2_, ∼310 mOsm, pH 7.2) with 0.001 TTX, 0 Mg^2+^, and 2 Ca^2+^. Cells were pre-incubated for at least 30 min with 10 μM L-689,560 (L-689, 15 mM stock in DMSO), 100 μM 7CK (100 mM stock in H_2_O), 2 µM SB203580 (4 mM stock in DMSO), 100 µM NG-Nitro-L-arginine (L-NNA, 200 mM stock in 0.25 N HCL), all from Tocris, or 10 µM MK2 inhibitor III (20 mM stock in DMSO) from Cayman Chemical, as indicated. Cells were pre-incubated for at least 60 min with 1 µM peptides (2 mM stock in H_2_O). Peptides L-TAT-GESV: NH2-GRKKRRQRRRYAGQWGESV-COOH and L-TAT-GASA: NH2-GRKKRRQRRRYAGQWGASA-COOH were obtained from GenicBio.

### HFU stimulus

High-frequency uncaging (HFU) consisted of 60 pulses (720 nm; ∼8-10 mW at the sample) of 2 ms duration at 2 Hz delivered in ACSF containing (in mM): 2 Ca^2+^, 0 Mg^2+^, 2.5 MNI-glutamate, and 0.001 TTX. The laser beam was parked at a point ∼0.5-1 μm from the spine head in the direction away from the dendrite.

### Image analysis

Estimated spine volume was measured from background-subtracted green fluorescence using the integrated pixel intensity of a boxed region surrounding the spine head, as previously described (Woods et al., 2011). All shown images are maximum projections of three-dimensional (3D) image stacks after applying a median filter (3 × 3) to the raw image data.

### Electrophysiology

Whole-cell recordings (V_hold_ = -65 mV; series resistances 20-40 MΩ) were obtained from visually identified CA1 pyramidal neurons in slice culture (14-18 DIV, depths of 10-50 µm) at 25 °C in ACSF containing in mM: 2 CaCl_2_, 1 MgCl_2_, 0.001 TTX, 2.5 MNI-glutamate. 10 μM L-689 or 100 μM 7CK were included as indicated. Recording pipettes (∼7 MΩ) were filled with cesium-based internal solution (in mM: 135 Cs-methanesulfonate, 10 Hepes, 10 Na_2_ phosphocreatine, 4 MgCl_2_, 4 Na_2_-ATP, 0.4 Na-GTP, 3 Na L-ascorbate, 0.2 Alexa 488, and ∼300 mOsm, ∼pH 7.25). For each cell, baseline uEPSCs were recorded (5-6 test pulses at 0.1 Hz, 720 nm, 1 ms duration, 8-10 mW at the sample) from two spines (2-12 μm apart) on secondary or tertiary basal branches (50-120 μm from the soma). The HFU stimulus was then applied to one spine, during which the cell was depolarized to 0 mV. Following the HFU stimulus, uEPSCs were recorded from both the target and neighboring spine at 5 min intervals for 25 min. uEPSC amplitudes from individual spines were quantified as the average from a 2 ms window centered on the maximum current amplitude within 50 ms following uncaging pulse.

### Statistics

All data are represented as mean ± standard error of the mean (SEM). All statistics were calculated across cells. Statistical significance was set at p < 0.05 (two-tailed t test). All statistical tests and p values are presented in **Supplemental Table 1**.

## Results

### p38 MAPK activity is required for spine shrinkage induced by non-ionotropic NMDAR signaling

In order to determine the signaling molecules downstream of non-ionotropic NMDAR function in spine shrinkage, we began by confirming a role for p38 MAPK, the only protein identified to date as a required component of the non-ionotropic NMDAR signaling cascade. We recently reported that p38 MAPK, which has been shown to play a role in conventional NMDAR-dependent LTD induced by low frequency stimulation (Zhu et al., 2002), is required downstream of non-ionotropic NMDAR signaling to drive dendritic spine shrinkage induced by a low-frequency uncaging stimulus (Stein et al., 2015) that also induces single spine LTD (Oh et al., 2013).

To assess the generalizability of the requirement for p38 MAPK as a signaling molecule downstream of metabotropic NMDAR function in spine shrinkage, we tested whether p38 MAPK was also required for spine shrinkage induced by high frequency uncaging (HFU, 60 pulses of 2 ms duration at 2 Hz) of glutamate in the presence of the NMDAR glycine/D-serine site antagonist 7-CK, which blocks ion flow through the NMDAR but leaves glutamate binding intact and, thus, converts a normally spine growth inducing HFU stimulus (**Fig. 1A-C**; veh: 216.4 ± 37.7%) into spine shrinkage. Indeed, we found that NMDAR-dependent spine shrinkage induced by HFU in the presence of 7-CK (**Fig. 1A-C**; 72.0 ± 5.1%) was blocked by the p38 MAPK inhibitor SB203580 (**Fig. 1A-C**; 114.3 ± 4.7%). Spine size of unstimulated neighboring spines was not affected (**Fig. 1A-C**; veh: 101.8 ± 6.1%; 7CK: 104.7 ± 2.2%; 7CK + SB: 101.3 ± 4.5%), excluding any acute independent effects of SB203580 on spine morphology. Thus, p38 MAPK is required for spine shrinkage driven by non-ionotropic NMDAR signaling in response to both low frequency and high frequency glutamatergic stimuli.

**Figure 1.**
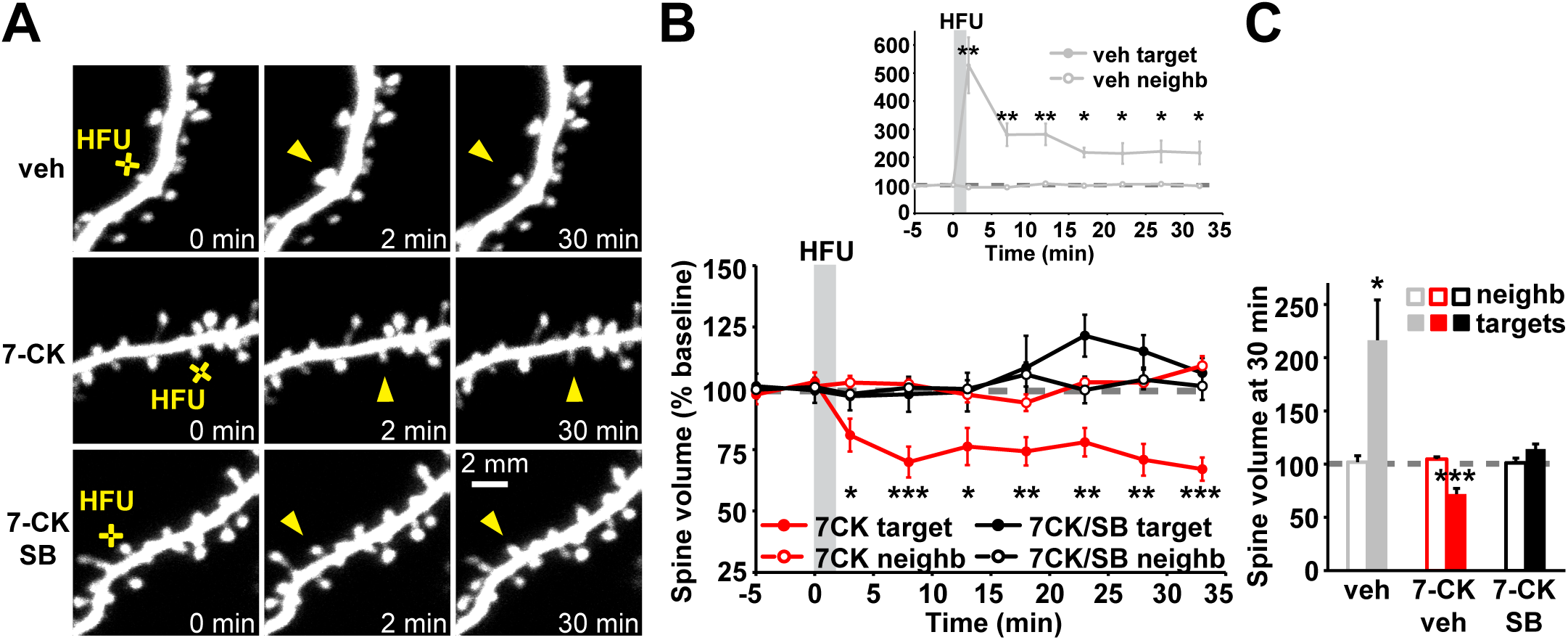
p38 MAPK activity is required for spine shrinkage driven by non-ionotropic NMDAR signaling in response to high frequency glutamate uncaging. **(A)** Images of dendrites from EGFP-transfected CA1 neurons at DIV14-18 before and after high-frequency glutamate uncaging (HFU, yellow cross) at individual dendritic spines (yellow arrowhead) in vehicle and in the presence of 7-CK (100 µM) with and without the p38 MAPK inhibitor SB203580 (SB, 2 µM). **(B, C)** HFU stimulation during vehicle conditions led to long-lasting spine growth (gray filled circles/bar; 6 spines/6 cells). However, in the presence of 7-CK the same uncaging stimulus induced dendritic spine shrinkage (red filled circles/bar; 11 spines/11 cells), which was blocked following inhibition of p38 MAPK activity with SB (black filled circles/bar; 8 spines/8 cells). Volume of unstimulated neighbors (open circles/bars) was unaffected. *p < 0.05; **p < 0.01, ***p < 0.001.

### p38 MAPK activity is required for LTD driven by non-ionotropic NMDAR signaling at individual dendritic spines

Our results confirm that p38 MAPK is required for the shrinkage and elimination of individual dendritic spines driven by non-ionotropic NMDAR signaling (Stein et al., 2015) and others have shown that p38 MAPK is activated by metabotropic NMDAR signaling in LTD (Nabavi et al., 2013). However, whether spine shrinkage induced by non-ionotropic NMDAR signaling at individual dendritic spines is associated with long-term depression of synaptic strength, and whether non-ionotropic NMDAR-LTD requires activation of p38 MAPK remains unknown.

To test whether non-ionotropic NMDAR-dependent signaling at individual dendritic spines also leads to LTD, we first needed to replace the glycine/D-serine site antagonist that we were using in our experiments because 7-CK potently inhibits AMPA receptor (AMPAR) currents (Leeson et al., 1992; Wong and Gray, 2018), making LTD experiments challenging. Compared to 7-CK, L-689,560 (L-689) is a more potent and selective glycine/D-serine site antagonist (Leeson et al., 1992), which completely blocks NMDAR-dependent ion flow at 10 μM (compared to 100 μM for 7-CK) and shows reduced inhibition of glutamate uncaging-evoked currents (uEPSCs) from AMPARs (**Fig. 2A**; 7-CK: 22.7 ± 2.8% of baseline; L-689: 64.4 ± 5.6% of baseline). Before initiating electrophysiological experiments, we first confirmed that L-689, like 7-CK, converted HFU-induced spine growth to spine shrinkage, characteristic of non-ionotropic NMDAR signaling. Indeed, HFU-induced spine shrinkage is observed in the presence of 10 μM L-689 (**Fig. 2B, C**; 57.3 ± 11.7%) similar to that found with 100 μM 7-CK (**Fig. 2B, C**; 58.6 ± 12.0%). Since 7-CK and L-689 allow partial AMPAR activation, we used 10 μM NBQX, a competitive AMPAR antagonist, to test whether metabotropic NMDAR signaling drives spine shrinkage entirely independent of AMPAR activation. Indeed, HFU-induced spine shrinkage in the presence of 100 μM 7-CK and 10 μM NBQX (**Fig. 2B, C**; 61.9 ± 11.3%) was not different than that observed with 7-CK or L-689 alone (p = 0.96).

**Figure 2.**
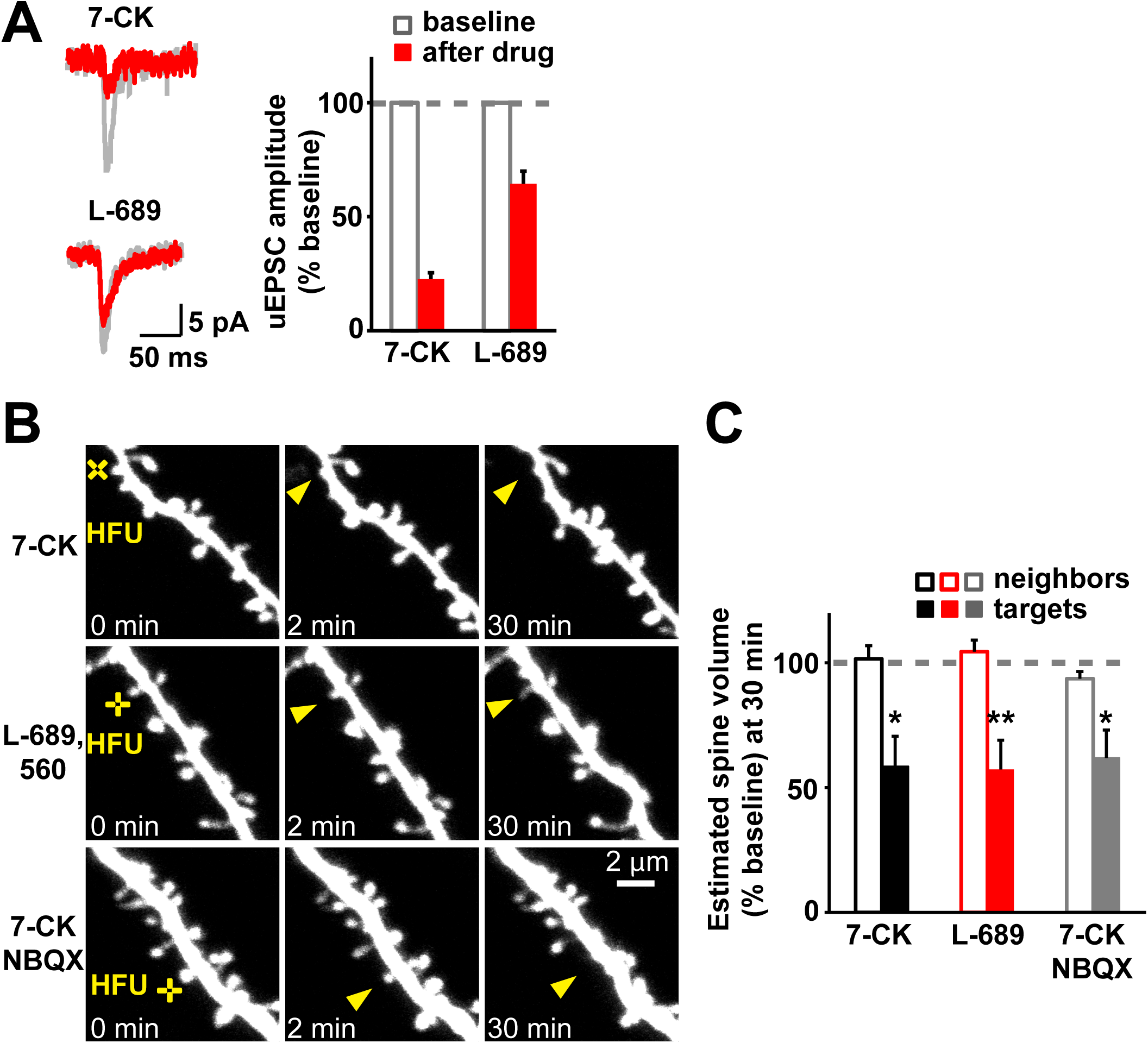
Spine shrinkage is induced by HFU in the presence of a lower concentration of L-689,560, which inhibits AMPARs to a lesser extent than 7-CK. **(A)** Left, representative uncaging-induced current traces (uEPSCs) from individual spines before (gray) and after (red) application of the NMDAR glycine/D-serine site antagonists 7-CK and L-689,560 (L-689). Right, compared to baseline (open gray bars) application of 100 µM 7-CK (22 spines/10 cells) greatly and 10 µM L-689 (6 spines/3 cells) partially reduced AMPAR uEPSCs (red filled bars). **(B)** Representative images of dendrites from EGFP-transfected CA1 neurons at DIV14-18 before and after HFU stimulation (yellow crosses) at individual dendritic spines (yellow arrowheads) in the presence of 100 µM 7CK, 10 µM L-689 or 100 µM 7-CK and 10 µM NBQX. **(C)** HFU stimulation in the presence of 7CK (black bar; 6 spines/6 cells), L-689 (red bar; 8 spines/8 cells) or a combination of 7CK and NBQX (gray bar; 7 spines/7 cells) caused a stable decrease in spine size at 30 min. Spine volume of the respective unstimulated neighbors (open bars) was not changed. *p < 0.05; **p < 0.01.

To test whether spine shrinkage induced by non-ionotropic NMDAR signaling at individual dendritic spines is associated with LTD, we recorded uEPSCs from one target spine and one neighboring spine at 5 min intervals before and after HFU stimulation in the presence of L-689. We found that HFU stimulation in the presence of L-689 led to a long-term decrease in the amplitude of uEPSCs (**Fig. 3A, B**; 76.4 ± 6.5%). This decrease in uEPSC amplitude was completely blocked by application of the p38 MAPK inhibitor SB203580 (**Fig. 3C, D**; 105.7 ± 6.0%). Non-ionotropic NMDAR-dependent LTD was specific to the stimulated target spine, as uEPSC amplitude of the unstimulated neighboring spines did not change (**Fig. 3A-D**; L-689: 105.7 ± 8.7%; L-689 + SB: 104.3 ± 7.8%). Together, our results indicate that p38 MAPK activity is required for both spine shrinkage and LTD induced by metabotropic NMDAR signaling.

**Figure 3.**
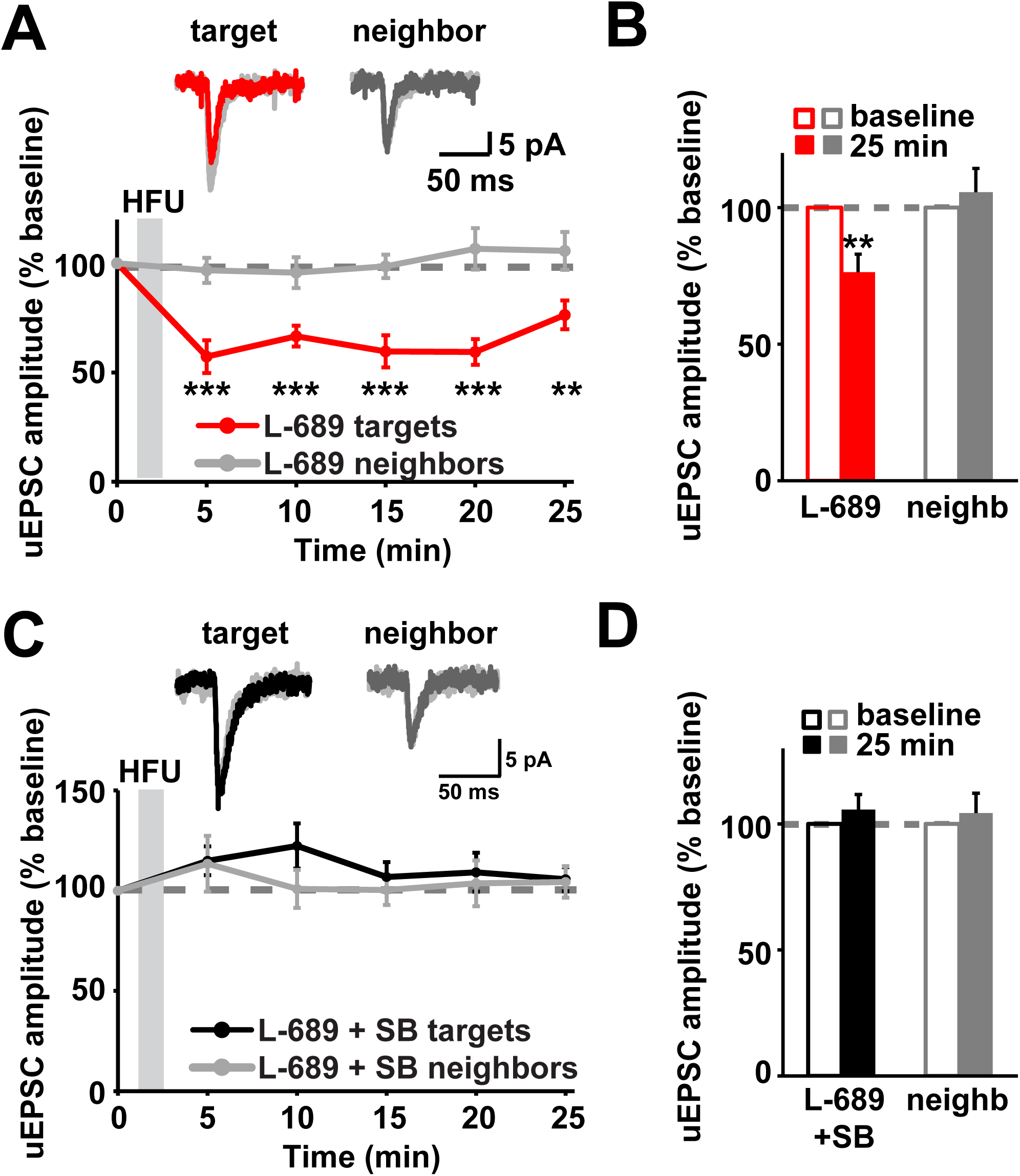
p38 MAPK activity is required for LTD driven by non-ionotropic signaling through the NMDAR. **(A)** Top, representative traces of uEPSCs from a target spine and an unstimulated neighbor before (light gray) and 25 min after HFU stimulation in the presence of L-689 (target, red; neighbor, dark gray). Bottom, averaged time course of uEPSC amplitude changes compared to baseline. **(B)** HFU stimulation in the presence of L-689 induced a long-lasting decrease in the uEPSC amplitude of stimulated spines (red line/bar; 7 spines/7 cells), while amplitude of unstimulated neighboring spines (gray line/bar) did not change. **(C)** Top, representative traces of uEPSCs from a target spine and an unstimulated neighbor before (gray) and 25 min after HFU stimulation in the presence of L-689 and the p38 MAPK inhibitor SB (target, red; neighbor, dark gray). Bottom, averaged time course of uEPSC amplitude changes compared to baseline.**(D)** The p38 MAPK inhibitor SB blocked LTD induced by HFU stimulation in the presence of L-689 (black line/bar; 7 spines/7 cells), while amplitude of unstimulated neighboring spines (gray line/bar) did not change. *p < 0.05; **p < 0.01, ***p < 0.001.

### NOS1AP interaction with nNOS, and nNOS enzymatic activity act downstream of conformational NMDAR signaling to drive dendritic spine shrinkage

In order to shed light on how p38 MAPK activation is driven by conformational changes of the NMDAR, we searched the literature for signaling proteins that could link the NMDAR to p38 MAPK activation. Intriguingly, Nitric Oxide Synthase 1 Adaptor Protein (NOS1AP) was recently implicated in p38 MAPK activation during NMDA-induced excitotoxicity (Li et al., 2013). Notably, selective disruption of the interaction between NOS1AP and nNOS with a cell permeant peptide L-TAT-GESV inhibited NMDAR-dependent p38 MAPK activation (Li et al., 2013; Li et al., 2015).

We tested whether L-TAT-GESV interferes with dendritic spine shrinkage driven by conformational signaling through the NMDAR. We found that application of L-TAT-GESV completely blocked spine shrinkage induced by HFU in the presence of 7-CK (**Fig. 4A-C**; 102.1 ± 9.1%), whereas the control peptide L-TAT-GASA, which does not compete with NOS1AP for the interaction with nNOS (Li et al., 2013), did not interfere with long-lasting dendritic spine shrinkage (**Fig. 4A-C**; 36.1 ± 6.9%). Importantly, neither the active peptide L-TAT-GESV or the control peptide L-TAT-GASA affected the volume of unstimulated neighboring spines (Fig. **4A-C**; 7CK + L-TAT-GASA: 93.5 ± 4.5%; 7CK + L-TAT-GESV: 107.7 ± 3.7%). Thus, interaction between nNOS and NOS1AP is required for spine shrinkage downstream of conformational NMDAR signaling.

**Figure 4.**
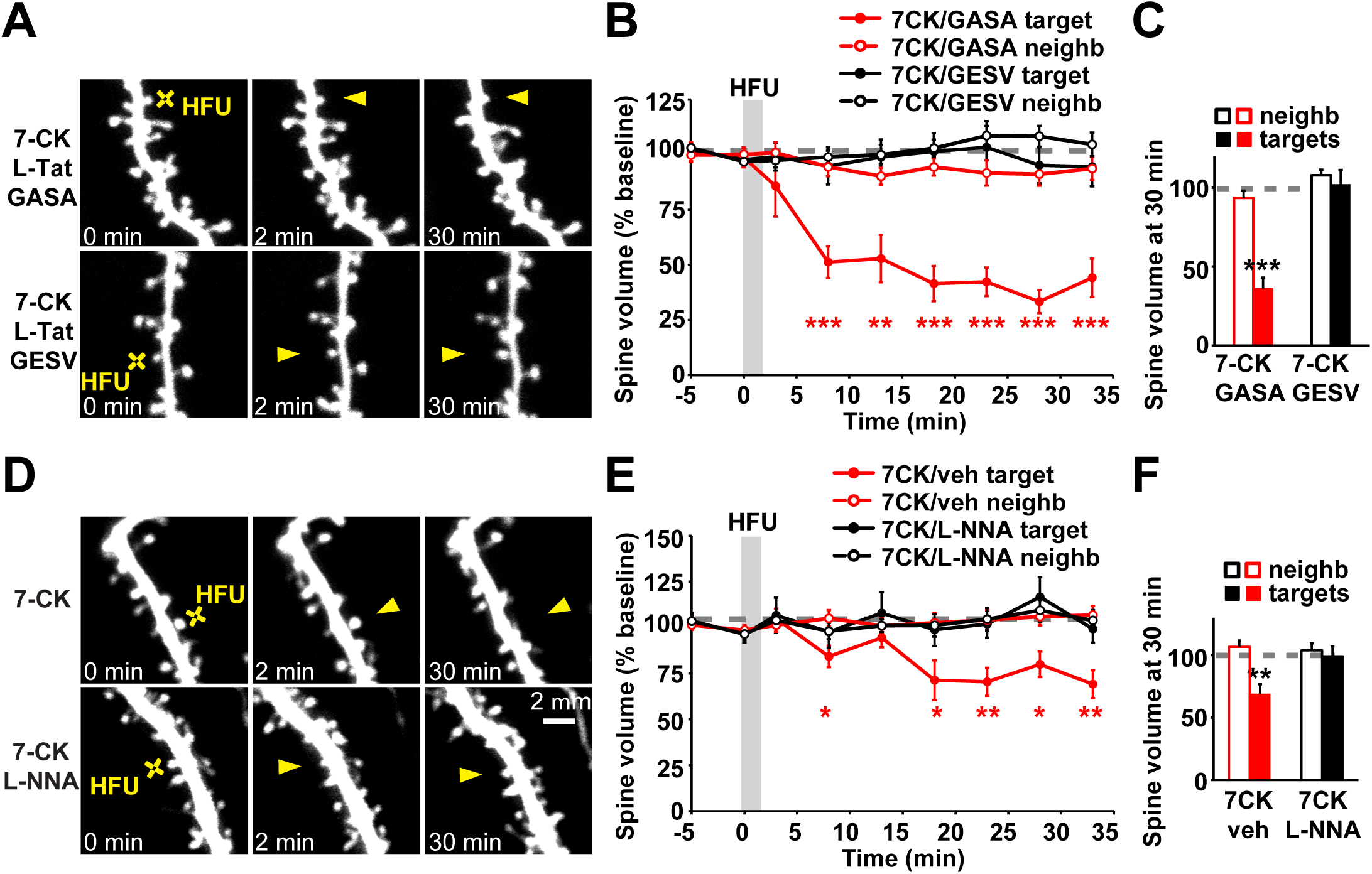
NOS1AP interaction with nNOS, and nNOS enzymatic activity are required for spine shrinkage driven by non-ionotropic signaling through the NMDAR. **(A)** Images of dendrites from EGFP-transfected CA1 neurons at DIV14-18 before and after high frequency glutamate uncaging (HFU, yellow cross) at an individual dendritic spine (yellow arrowhead) in the presence of 7CK (100 µM) and L-TAT-GESV (1 µM) or L-TAT-GASA (1 µM). **(B, C)** Disruption of NOS1AP/nNOS interaction using the active cell permeant L-TAT-GESV peptide (black filled circles/bar; 15 spines/15 cells), but not the inactive L-TAT-GASA control peptide (red filled circles/bar; 8 spines/8 cells), blocked spine shrinkage induced by non-ionotropic NMDAR signaling. Spine volume of the unstimulated neighbors (open circles/bars) was unchanged. **(D)** Images of dendrites from EGFP-transfected CA1 neurons at DIV14-18 before and after high frequency glutamate uncaging (HFU, yellow cross) at an individual dendritic spine (yellow arrowhead) in the presence of 7-CK (100 µM) or 7-CK (100 µM) and L-NNA (100 µM). **(E, F)** Inhibition of NO synthase activity with L-NNA blocked spine shrinkage (solid black circles/bar; 11 spines/11 cells) induced by HFU (yellow cross) in the presence of 7-CK (solid red circles/bar; 13 spines/13 cells). Spine volume of the unstimulated neighbors (open circles/bars) was unchanged. **p < 0.01, ***p < 0.001.

It is possible that nNOS simply functions as a scaffolding molecule to recruit NOS1AP into the NMDAR complex via its interactions with PSD-95 (Christopherson et al., 1999). Alternatively, nNOS enzymatic activity might be required for metabotropic NMDAR signaling. We tested whether nNOS enzymatic activity is required using the NO synthase inhibitor NG-Nitro-L-arginine (L-NNA). We found that application of L-NNA abolished HFU-induced non-ionotropic NMDAR-dependent spine shrinkage (**Fig. 4D-F**; 7-CK: 69.2 ± 7.6%; 7-CK + L-NNA: 99.3 ± 7.6%). Importantly, L-NNA did not affect the volume of unstimulated neighboring spines (**Fig. 4D-F**; 7CK: 105.7 ± 5.1%; 7-CK + L-NNA: 103.9 ± 5.7%). Together, our results support a model where non-ionotropic NMDAR function drives dendritic spine shrinkage through nNOS activity and its interaction with NOS1AP.

### MK2 activity and cofilin are required downstream of conformational NMDAR signaling to drive dendritic spine shrinkage

Spine shrinkage following LTD induction relies upon remodeling of the spine actin cytoskeleton through the action of the actin depolymerizing protein cofilin (Zhou et al., 2004; Wang et al., 2007; Hayama et al., 2013). To shed light on how non-ionotropic NMDAR signaling leads to remodeling of the actin cytoskeleton in spine shrinkage, we searched for signaling proteins that could link p38 MAPK to cofilin.

Interestingly, during mGluR-dependent LTD, a role for p38 MAPK and its downstream substrate MAPK activated protein kinase 2 (MK2) was identified in the regulation of cofilin activity and dendritic spine morphology (Eales et al., 2014). We tested whether spine shrinkage driven by metabotropic NMDAR signaling is dependent on MK2 activity. We found that spine shrinkage induced by HFU in the presence of 7-CK was blocked by application of MK2 inhibitor III (**Fig. 5A-C**; 7-CK: 70.5 ± 6.4%; 7-CK + MK2 inhibitor III: 105.5 ± 11.7%). Importantly, MK2 inhibitor III did not affect the size of unstimulated neighboring spines (**Fig. 5A-C**; 7-CK: 93.9 ± 3.9%; 7-CK + MK2 inhibitor III: 106.0 ± 4.2%). Thus, MK2 activity is required for spine shrinkage driven by non-ionotropic NMDAR signaling.

**Figure 5.**
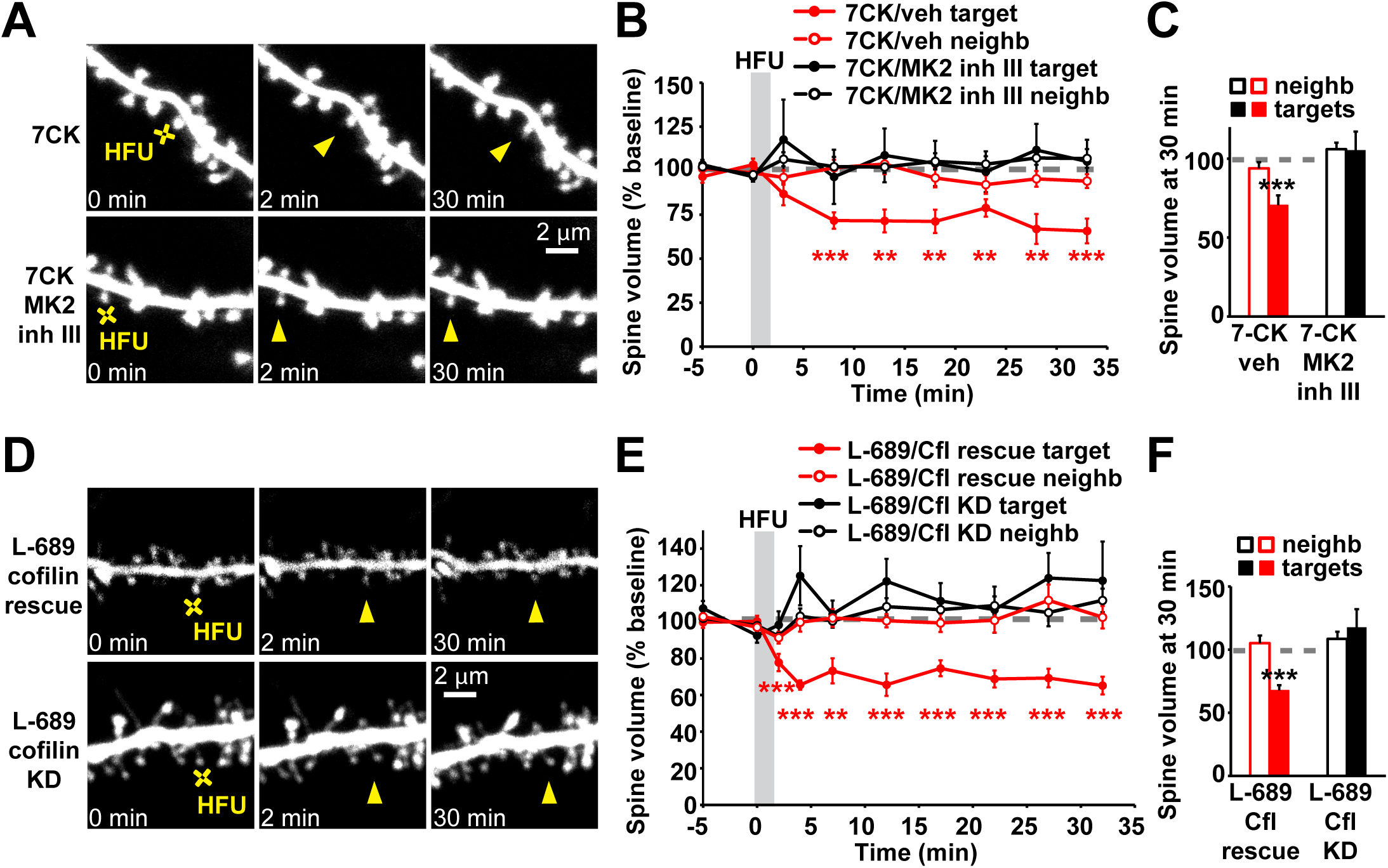
MK2 activity and cofilin are required for spine shrinkage driven by non-ionotropic NMDAR signaling. **(A)** Images of dendrites from EGFP-expressing neurons (DIV14-18) showing 7-CK-dependent HFU-induced (yellow crosses) spine shrinkage (yellow arrowheads) in the presence of MK2 inhibitor III (10 µM). **(B, C)** Inhibition of MK2 activity (black filled circles/bar; 11 spines/11 cells) prevented 7-CK dependent non-ionotropic spine shrinkage (red filled circles/bar; 10 spines/10 cells). Spine volume of the unstimulated neighbors (open circles/bars) did not change. **(D)** Images of dendrites from DIV14-18 neurons expressing cofilin and ADF shRNAs (KD) together with EGFP and CyRFP1 or in combination with shRNA-resistant cofilin-EGFP and CyRFP1 before and after HFU stimulation (yellow crosses) at a single dendritic spine (yellow arrowheads) in the presence of L-689 (10 µM). **(E, F)** KD of cofilin and ADF (black filled circles/bar; 11 spines/11 cells) blocked non-ionotropic NMDAR-dependent spine shrinkage in the presence of L-689 and was rescued by shRNA-resistant cofilin-EGFP expression (red filled circles/bar; 9 spines/9 cells). Spine volume of the respective unstimulated neighbors (open circles/bars) was not changed. *p < 0.05; **p < 0.01, ***p < 0.001.

Furthermore, to confirm that cofilin is required downstream of this non-ionotropic NMDAR- and p38 MAPK-dependent signaling in spine shrinkage, we knocked down cofilin together with actin depolymerizing factor (ADF, a member of the cofilin protein family) using previously published shRNA constructs (Bosch et al., 2014). We found that knock down of cofilin and ADF blocked HFU-induced dendritic spine shrinkage in the presence of L-689 (**Fig. 5D-F**; shRNAs/L-689: 117.7 ± 14.3%). Spine shrinkage was restored by co-expression of a shRNA-resistant version of wild-type cofilin (**Fig. 5D-F**; shRNAs + cofilin rescue/L-689: 67.8 ± 4.1%). Importantly, spine size of unstimulated neighbors was not changed in either case (**Fig. 5D-F**; shRNAs/L-689: 108.6 ± 5.6%; shRNAs + cofilin rescue/L-689: 105.1 ± 6.1%). Thus, cofilin activation is required for spine shrinkage induced by metabotropic NMDAR signaling.

We further investigated the role of cofilin by monitoring the redistribution of cofilin-GFP following induction of structural plasticity by HFU stimulation. Using cells co-expressing cofilin-GFP and the red cell fill CyRFP1, we simultaneously monitored changes in cofilin-GFP and spine volume. We found that there was no change in cofilin-GFP levels relative to spine volume immediately following HFU stimulation in the presence of 7-CK, but 10 min after uncaging the amount of cofilin-GFP decreased compared to spine volume and stayed decreased for at least up to 30 min (**Fig. 6A,B**; cofilin-GFP/CyRFP1 ratio at 25-35 min following HFU stimulation: 85.7 ± 4.3%). As a control, we show that during HFU-induced structural LTP, cofilin-GFP enriches in the stimulated spine for at least up to 30 min (**Fig. 6C,D**; cofilin-GFP/CyRFP1 ratio at 25-35 min following HFU stimulation: 217.1 ± 43.3%) as previously reported (Bosch et al., 2014).

**Figure 6.**
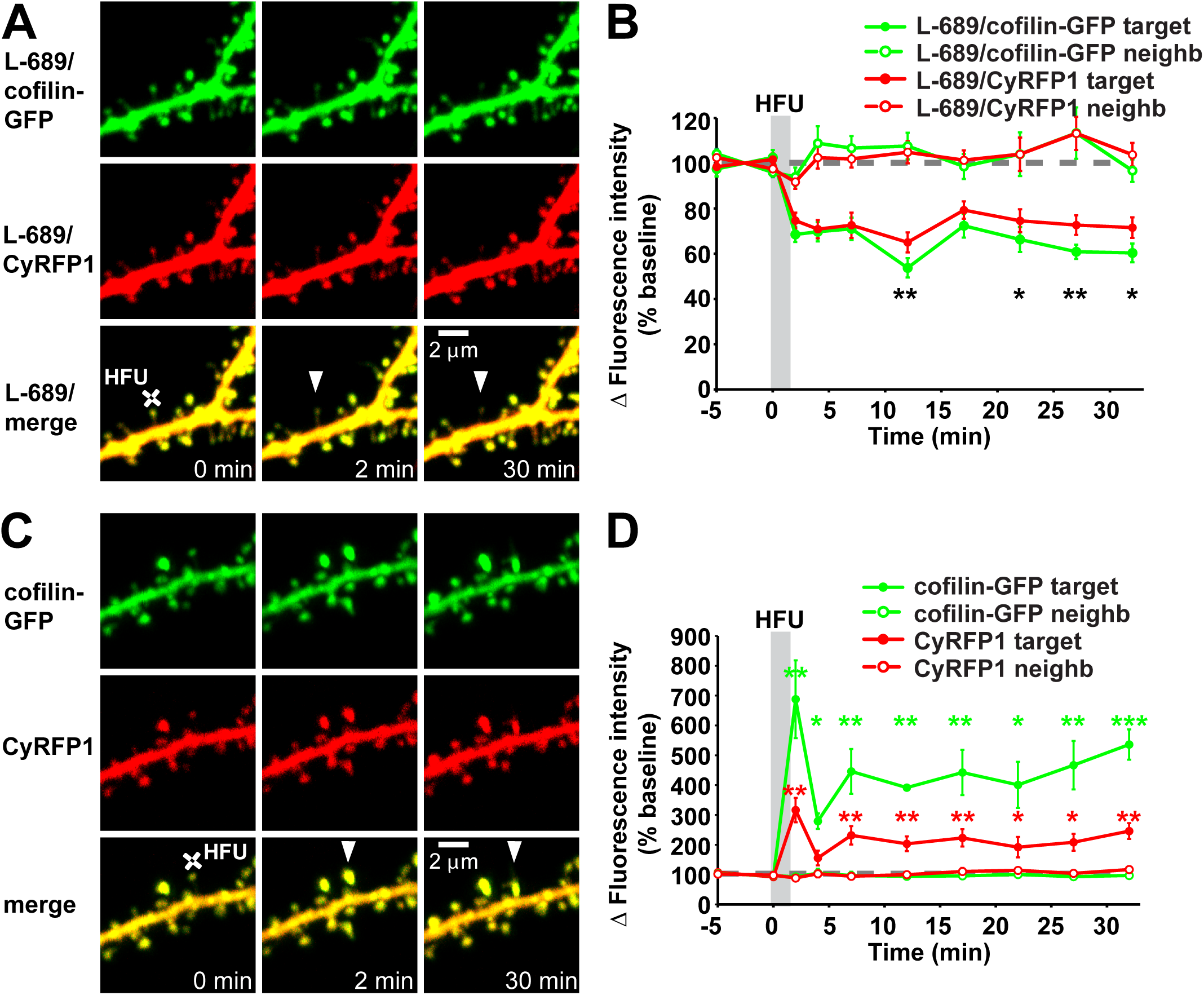
Non-ionotropic NMDAR-dependent spine shrinkage is associated with loss of cofilin from the spine head. **(A)** Images of CA1 neurons transfected with cofilin shRNAs in combination with shRNA-resistant cofilin-EGFP and CyRFP1 before and after HFU stimulation (white crosses) of individual spines (white arrowheads) in the presence of L-689. **(B)** Spine shrinkage (red, CyRFP1) induced by HFU in the presence of L-689 was associated with a decrease of cofilin-GFP protein levels in the spine (green) compared to baseline. Cofilin-GFP spine levels were decreased 10 min after HFU in L-689 and stayed decreased until 30 min after HFU in L-689 (16 spines/16 cells). **(C)** Images of dendrites from DIV14-18 CA1 neurons expressing cofilin shRNAs in combination with shRNA-resistant cofilin-EGFP and CyRFP1 before and after HFU stimulation (white crosses) of individual spines (white arrowheads). **(D)** Time course showing HFU-induced changes in spine volume (red, CyRFP1) and the amount of cofilin-GFP protein in the spine (green) compared to baseline. Cofilin-GFP spine levels were enriched after sLTP induction and remained enriched for at least 30 min following HFU (6 spines/6 cells). *p < 0.05; **p < 0.01, ***p < 0.001.

### CaMKII activity is required for spine shrinkage driven by conformational NMDAR signaling

CaMKII has been shown to reposition within the NMDAR complex in response to metabotropic NMDAR signaling (Aow et al., 2015), suggesting that it might play a role in signaling downstream of conformational signaling through the NMDAR. Notably, CaMKII, which has been extensively studied in LTP induction (Lisman et al., 2012; Hell, 2014), lately also has been implicated in LTD (Coultrap et al., 2014; Goodell et al., 2017; Woolfrey et al., 2018), further supporting a possible role in spine shrinkage downstream of non-ionotropic NMDAR signaling.

We tested whether CaMKII is required for spine shrinkage driven by conformational NMDAR signaling. We found that dendritic spine shrinkage induced by HFU in the presence of L-689 (**Fig. 7A-C**; L-689: 75.4 ± 3.7%) was blocked in the presence of KN-62 (**Fig. 7A-C**; L-689 + KN-62: 92.9 ± 4.4%). Importantly, size of unstimulated neighboring spines was not affected (**Fig. 7A-C**; L-689: 95.3 ± 5.0%; L-689 + KN-62: 90.3 ± 2.4%). Our results demonstrate that CaMKII is required for dendritic spine shrinkage induced by conformational NMDAR signaling.

**Figure 7.**
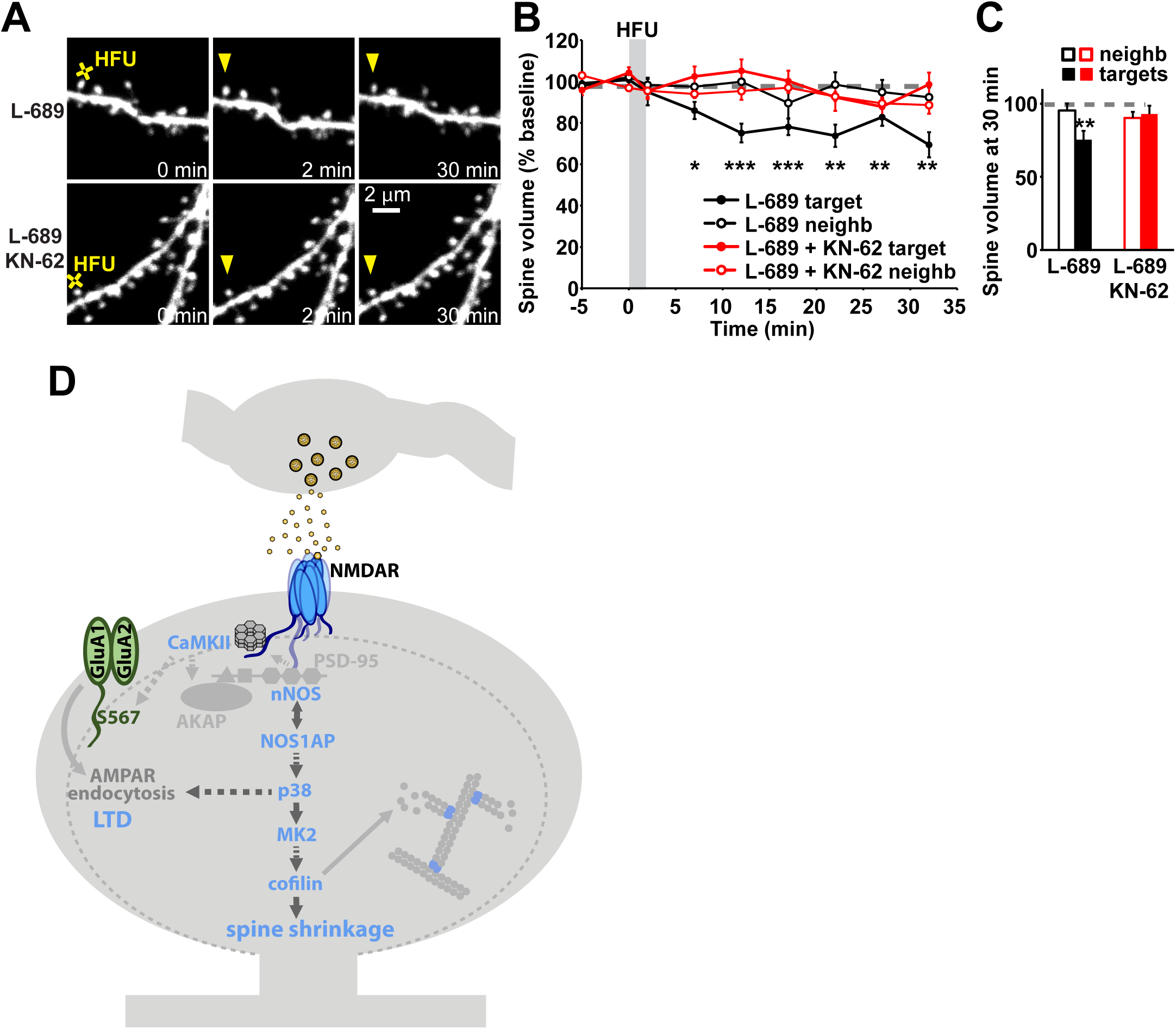
CaMKII activity is required for spine shrinkage driven by non-ionotropic NMDAR signaling. **(A)** Images of dendrites from CA1 neurons of acute slices from P16-20 GFP-M mice before and after HFU stimulation (yellow cross) of single spines (yellow arrowhead) in the presence of L-689 or L-689 with KN-62. **(B, C)** Inhibition of CaMKII activity with KN-62 (red filled circles/bar; 9 spines/9 cells) blocked non-ionotropic NMDAR-dependent spine shrinkage (black filled circles/bar; 8 spines/8 cells). Spine volume of the respective unstimulated neighbors (open bars) was not changed. *p < 0.05; **p < 0.01, ***p < 0.001. **(D)** Proposed model for the non-ionotropic NMDAR signaling pathway that drives spine shrinkage. Glutamate binding to the NMDAR induces conformational changes that, in the absence of ion influx through the NMDAR, drive dendritic spine shrinkage through NOS1AP-nNOS interactions, the activities of nNOS, p38 MAPK, MK2, CaMKII, and cofilin-dependent severing of the actin cytoskeleton.

## Discussion

### Molecular mechanisms of non-ionotropic NMDAR signaling

Despite several recent studies demonstrating that the NMDAR can signal independent of ion flow to drive dendritic spine shrinkage and LTD (Nabavi et al., 2013; Aow et al., 2015; Stein et al., 2015; Carter and Jahr, 2016; Wong and Gray, 2018), the molecular signaling mechanisms that link conformational NMDAR signaling to LTD and spine shrinkage remained poorly defined. Here, we have identified several components of key importance in this signaling cascade.

### p38 MAPK

Only one protein, p38 MAPK, had been implicated as a downstream component of non-ionotropic signaling by the NMDAR. p38 MAPK was shown to be activated downstream of metabotropic NMDAR function in LTD (Nabavi et al., 2013) and to be required for spine shrinkage driven by metabotropic NMDAR signaling (Stein et al., 2015), both of which were induced by low frequency glutamatergic stimulation. Here, we identify p38 MAPK as a central signaling component in additional paradigms of non-ionotropic NMDAR signaling during synaptic plasticity. First, we show that p38 MAPK is required for spine shrinkage induced by high frequency glutamatergic stimulation in the presence of the glycine/D-serine site NMDAR antagonists, 7-CK and L-689. Second, we show that p38 MAPK is required for long-term depression of synaptic currents at individual dendritic spines induced by high frequency glutamatergic stimulation in the presence of the glycine/D-serine site NMDAR antagonist, L-689. Combined, these results confirm a key role for p38 MAPK in the signaling cascade driven by conformational signaling by the NMDAR.

### NOS1AP and nNOS

We identified a novel role for the interaction between NOS1AP and nNOS, likely upstream of p38 MAPK, in spine shrinkage induced by non-ionotropic NMDAR signaling. We propose that the recruitment of NOS1AP to nNOS is important for the localized activation of p38 MAPK and allows for the subsequent phosphorylation of LTD specific targets driving spine shrinkage and AMPAR endocytosis. Indeed, it has been previously shown that NOS1AP interacts with the p38 MAPK activator MKK3 and that during NMDA-induced excitotoxicity NOS1AP is recruited to nNOS (Li et al., 2013). Furthermore, p38 MAPK activation is dependent on both the NOS1AP/nNOS interaction and MKK3 (Li et al., 2013). Linked to PSD-95 and the NMDAR complex, p38 MAPK could participate with Rap1 in microdomain specific signaling at late endosomes and could contribute to S880 phosphorylation of GluA2, which disrupts AMPAR anchoring (Chung et al., 2000; Zhang et al., 2018a), leading to LTD and spine shrinkage.

In addition, it has been shown that nNOS activity is required for NOS1AP recruitment during excitotoxicity (Li et al., 2013) and that the interaction with NOS1AP also is important for nNOS-mediated nitrosylation of Dexras1, which negatively regulates Erk (Zhu et al., 2014; Zhang et al., 2018b). Thus, this dual and opposite regulation of p38 and Erk MAPK activity through NOS1AP, if occurring during non-ionotropic NMDAR signaling, would allow NOS1AP to locally activate p38 MAPK-dependent LTD and spine shrinkage signaling pathways and at the same time downregulate Erk-dependent LTP signaling (Zhu et al., 2002; Zhang et al., 2018a). Alternatively, the effects of Dexras1 on Erk activity could be independent of the fast activity-induced recruitment of NOS1AP and upregulation of p38 MAPK activity as changes in Erk activity only were reported after long-term treatments.

### MK2 and cofilin

Downstream of glutamate-induced conformational changes in the NMDAR (Dore et al., 2015; Ferreira et al., 2017), we showed that the p38 MAPK substrate, MAPK activated protein kinase 2 (MK2), and cofilin are required for non-ionotropic NMDAR-dependent spine shrinkage. Earlier studies showed that MK2 is activated by p38 MAPK during DHPG-induced mGluR dependent LTD and that mGluR-dependent LTD leads to dephosphorylation and activation of cofilin in a p38 MAPK-dependent manner (Eales et al., 2014).

We propose that conformational NMDAR signaling also leads to activated p38 MAPK that will bind and phosphorylate MK2, and that active MK2 is important for activation of cofilin. Yet the link between activation of MK2 and the decrease in cofilin1 phosphorylation remains unclear. The severing and depolymerization activity of cofilin is regulated by phosphorylation on Ser3 through an interplay of the deactivating kinase LIMK1 and the phosphatase slingshot 1 (SSH1). In endothelial cells MK2 has been shown to phosphorylate and activate LIMK1 on a secondary extracatalytic site (S323), next to the classical Rho family small GTPase-dependent phosphorylation and activation of LIMK1 in the catalytic domain (S508) (Kobayashi et al., 2006; Scott and Olson, 2007). Increases in LIMK1 phosphorylation caused by MK2 activation and the subsequent F-actin stabilization could activate slingshot 1, which consequently dephosphorylates LIMK1 and cofilin.

Alternatively, MK2 could act on a target independent of LIMK1, as observed during bone morphogenetic protein-2 induced cell migration, where the p38 MAPK-MK2-small heat-shock protein 25 cascade is required for actin remodeling (Gamell et al., 2011). Further complicating the exact regulation of cofilin is the fact that, in contrast to LIMK1, not much is known about synapse specific pathways regulating slingshot activity and other phosphatases like PP1 have been shown to be able to activate cofilin (Ohashi, 2015; Shaw and Bamburg, 2017). In addition, PP2B, which at basal activity levels also is required for non-ionotropic signaling, has been shown to be required together with slingshot 1 for F-actin reorganization during ephrin A induced spine retraction (Zhou et al., 2012).

### CaMKII

We show that CaMKII activity is required for spine shrinkage driven by non-ionotropic NMDAR signaling. This may appear surprising, as the majority of studies on CaMKII have focused on its role in LTP and spine growth; however, several recent studies have revealed a role of CaMKII in LTD (Coultrap et al., 2014; Goodell et al., 2017; Woolfrey et al., 2018), consistent with our finding of a role for CaMKII in spine shrinkage.

We propose that the NMDAR conformational change could move NMDAR-bound CaMKII closer to different substrates. CaMKII would normally be out of reach of these targets, and be brought in close proximity with non-ionotropic NMDAR conformational change. Indeed, in the LTD studies, autonomously active CaMKII was shown to have different substrate selectivity than when Ca^2+^/CaM is bound, targeting non-traditional substrates like AKAP79/150 or GluA1 S567 to mediate LTD (Coultrap et al., 2014; Woolfrey et al., 2018). However, in contrast, it has been observed that CaMKII bound to the NMDAR exhibits decreased autonomous activity, which is accompanied by a delayed repositioning within the receptor complex (Aow et al., 2015). We expect that CaMKII repositioning within the NMDAR complex and T286 dephosphorylation may occur as a safeguard to limit autonomous CaMKII activity during non-ionotropic NMDAR signaling.

Alternatively, or additionally, CaMKII may have a structural role in the non-ionotropic NMDAR signaling pathway through its interaction with F-actin. CaMKII-actin interaction is known to stabilize F-actin by crosslinking the actin filaments and preventing actin-regulating proteins from binding to F-actin (Hell, 2014; Kim et al., 2015). Release of CaMKII-actin interaction is crucial for both LTP and spine growth by allowing actin-regulating proteins like cofilin to bind to F-actin (Kim et al., 2015). Because non-ionotropic NMDAR signaling requires cofilin activity to drive spine shrinkage, we expect that CaMKII-actin interaction must be released as well.

### Non-ionotropic NMDAR signaling in disease

Here, we identified a novel role for the interaction of NOS1AP and nNOS in spine shrinkage driven by non-ionotropic NMDAR signaling. Notably, both nNOS and NOS1AP have been identified as schizophrenia risk genes (Shinkai et al., 2002; Freudenberg et al., 2015). Our findings raise the question whether NOS1AP-mediated signaling contributes to the spine loss associated with schizophrenia. Intriguingly, reduced levels of the synaptic NMDAR co-agonist D-serine and polymorphisms of genes involved in the regulation of D-serine levels have been found in schizophrenic patients (Hashimoto et al., 2005; Goltsov et al., 2006; Balu et al., 2013). These pathologic conditions could result in a shift towards increased non-ionotropic NMDAR signaling and could contribute to the decreased spine density observed in patients with schizophrenia (Penzes et al., 2011; Glausier and Lewis, 2013).

Several earlier studies have implicated non-ionotropic NMDAR signaling in Alzheimer’s disease (Kessels et al., 2013; Tamburri et al., 2013; Birnbaum et al., 2015), which is associated with dendritic spine loss (Selkoe, 2002). Notably, p38 MAPK activity was linked to amyloid beta (Aβ)-induced spine loss driven by non-ionotropic NMDAR signaling (Birnbaum et al., 2015). In addition, in models of Alzheimer’s disease, increased nNOS-NOS1AP interaction was detected after treatment with Aβ in vitro and in APP/PS1 mice in vivo (Zhang et al., 2018b). After blocking the nNOS-NOS1AP interaction, memory was rescued in 4-month-old APP/PS1 mice, and dendritic impairments were ameliorated both in vivo and in vitro (Zhang et al., 2018b), further supporting a possible role for non-ionotropic NMDAR signaling in Alzheimer’s disease.

Thus, delineating the non-ionotropic NMDAR signaling pathway not only leads to a better understanding of the molecular mechanisms underlying structural plasticity of dendritic spines during experience-dependent plasticity, but also provides new insights into how dysregulation of these mechanisms could contribute to altered dendritic spine dynamics and density in neuropsychiatric and neurodegenerative diseases.

## Acknowledgements

This work was supported by the National Institutes of Health (R01 NS062736 to K.Z.). We thank Dr. Yasunori Hayashi for the cofilin related DNA constructs; Julie Culp and Lorenzo Tom for technical support and help with analysis, and Nicole Claiborne, Jinyoung Jang, and Samuel Petshow for critical reading of the manuscript.

## Figure Legends

**Supplemental Table 1.**

| **Figure** | **Sample Size** | **Analysis** | **P Values** |
| --- | --- | --- | --- |
| Fig. 1B, C | Vehicle: n = 6 stimulated spines from 6 different cells (6 spines/6 cells)<br>7CK + vehicle: n = 11 spines/11 cells<br>7CK + SB203580: n = 8 spines/8 cells<br>Cells for each condition were obtained from at least 3 different dissections from P6-P8 SD rats of both sexes. | Paired two-tailed T-Test over spines/cells | T(2, 7, 12, 17, 22, 27, 32 min)<br>P=0.008, 0.007, 0.005, 0.001, 0.027, 0.027, 0.035<br>T(2, 7, 12, 17, 22, 27, 32 min)<br>P=0.017, 0.001, 0.011, 0.001, 0.003, 0.001, 0.0001<br>T(2, 7, 12, 17, 22, 27, 32 min)<br>P=0.597, 0.744, 0.851, 0.538, 0.040, 0.056, 0.338 |
| Fig. 2C | 7CK: n = 6 spines/6 cells<br>L-689,560: n = 8 spines/8 cells<br>7CK + NBQX: n = 7 spines/7 cells<br>Cells for each condition were obtained from at least 3 different dissections from P6-P8 SD rats of both sexes. | Paired two-tailed T-Test over spines/cells | P=0.013<br>P=0.005<br>P=0.013 |
| Fig. 3A, B | L-689,560: n = 7 spines/7 cells<br>Cells were obtained from at least 3 different dissections from P6-P8 SD rats of both sexes. | Paired two-tailed T-Test over spines/cells | T(5, 10, 15, 20, 25 min)<br>P=0.001, 0.0002, 0.001, 0.0002, 0.009 |
| Fig. 3C, D | L-689,560 + SB203580: n = 7 spines/7 cells<br>Cells were obtained from at least 3 different dissections from P6-P8 SD rats of both sexes. | Paired two-tailed T-Test over spines/cells | T(5, 10, 15, 20, 25 min)<br>P=0.084, 0.095, 0.391, 0.381, 0.383 |
| Fig. 4B, C | 7CK + L-TAT-GASA: n = 8 spines/8 cells | Paired two-tailed T-Test over spines/cells | T(2, 7, 12, 17, 22, 27, 32 min)<br>P=0.166, 0.001, 0.002, 0.0002, 0.0001, 0.0001, 0.0003 |
|  | 7CK + L-TAT-GESV: n = 15 spines/15 cells<br>Cells for each condition were obtained from at least 3 different dissections from P6-P8 SD rats of both sexes. |  | T(2, 7, 12, 17, 22, 27, 32 min)<br>P=0.884, 0.955, 0.968, 0.848, 0.489, 0.689, 0.944 |
| Fig. 4E, F | 7CK + vehicle: n = 13 spines/13 cells<br>7CK + L-NNA: n = 11 spines/11 cells<br>Cells for each condition were obtained from at least 3 different dissections from P6-P8 SD rats of both sexes. | Paired two-tailed T-Test over spines/cells | T(2, 7, 12, 17, 22, 27, 32 min)<br>P=0.379, 0.020, 0.312, 0.022, 0.002, 0.014, 0.002<br>T(2, 7, 12, 17, 22, 27, 32 min)<br>P=0.502, 0.791, 0.502, 0.896, 0.803, 0.154, 0.928 |
| Fig. 5B, C | 7CK + vehicle: n = 10 spines/10 cells<br>7CK + MK2 inhibitor III: n = 11 spines/11 cells<br>Cells for each condition were obtained from at least 3 different dissections from P6-P8 SD rats of both sexes. | Paired two-tailed T-Test over spines/cells | T(2, 7, 12, 17, 22, 27, 32 min)<br>P=0.069, 0.0002, 0.002, 0.002, 0.002, 0.004, 0.001<br>T(2, 7, 12, 17, 22, 27, 32 min)<br>P=0.454, 0.825, 0.581, 0.759, 0.962, 0.447, 0.680 |
| Fig. 5E, F | L-689,560 + cofilin/ADF KD + cofilin rescue: n = 9 spines/9 cells<br>L-689,560 + cofilin/ADF KD: n = 11 spines/11 cells<br>Cells for each condition were obtained from at least 3 different dissections from P6-P8 C57BL/6J mice of both sexes. | Paired two-tailed T-Test over spines/cells | T(2, 4, 7, 12, 17, 22, 27, 32 min) P=0.001, 0.0001, 0.003, 0.0002, 0.0002, 0.0001, 0.0001, 0.0001<br>T(2, 4, 7, 12, 17, 22, 27, 32 min) P= 0.831, 0.157, 0.579, 0.107, 0.277, 0.612, 0.118, 0.322 |
| Fig. 6B | L-689,560 + cofilin KD + cofilin rescue: n = 16 spines/16 cells | Paired two-tailed T-Test over spines/cells | T(2, 4, 7, 12, 17, 22, 27, 32 min) P=0.043, 0.662, 0.623, 0.005, 0.087, 0.035, 0.004, 0.019 |
|  | Cells were obtained from at least 3 different dissections from P6-P8 C57BL/6J mice of both sexes. |  |  |
| Fig. 6C | vehicle + cofilin KD + cofilin rescue: n = 6 spines/6 cells<br>Cells were obtained from at least 3 different dissections from P6-P8 C57BL/6J mice of both sexes. | Paired two-tailed T-Test over spines/cells | T(2, 4, 7, 12, 17, 22, 27, 32 min) P=0.014, 0.136, 0.021, 0.018, 0.037, 0.023, 0.022, 0.008 |
| Fig. 7B, C | L-689,560 + vehicle: n = 8 spines/8 cells<br>L-689,560 + KN-62: n = 9 spines/9 cells<br>Cells for each condition were obtained from at least 4 different acute slice preparations from P16-P20 GFP-M mice of both sexes. | Paired two-tailed T-Test over spines/cells | T(2, 7, 12, 17, 22, 27, 32 min) P=0.456, 0.012, 0.001, 0.001, 0.002, 0.005, 0.002<br>T(2, 7, 12, 17, 22, 27, 32 min) P=0.286, 0.620, 0.363, 0.960, 0.279, 0.053, 0.815 |

